# Big Data, Sound Science, Lasting Impact: a framework for passive acoustic monitoring

**DOI:** 10.1101/2025.04.22.650104

**Authors:** Carrie C. Wall, Megan McKenna, Leila T. Hatch, Sofie M. Van Parijs, Rob Bochenek, Peter Dugan, Clea Parcerisas, John Ryan, Charles D. Anderson, Kyle Becker, Catherine Berchok, Mathew Biddle, Olaf Boebel, Adrienne Canino, Gabrielle Canonico, Danelle Cline, Genevieve E. Davis, Kaitlin Frasier, Jason Gedamke, Samara M. Haver, Karina Khazmutdinova, Niels Kinneging, Anurag Kumar, Alyssa Marian, Jennifer Miksis-Olds, Eric W. Montie, Dimitri Ponirakis, Aaron N. Rice, Timothy J. Rowell, Carlos Rueda, Emily Shumchenia, Thomas Shyka, Erica Staaterman, Karolin Thomisch

## Abstract

Marine passive acoustic monitoring (PAM) has produced petabytes of data that are used by researchers, resource managers, industry, and regulators to understand how marine animals use sound and the impacts of anthropogenic noise on species and ecosystems throughout the global ocean. These big data provide unprecedented opportunities to study underwater soundscapes and marine ecology but also enormous challenges to efficiently extract information. To address these challenges, a U.S. federally funded and led Sound Cooperative (*SoundCoop*) project built community-focused cyberinfrastructure to promote improved, scalable and sustainable processing and access of marine PAM data for management, science, industry and military applications. Driven by cross-institutional participation representing a diversity of data collection methods and conditions, the *SoundCoop* project established guidance for standardized processing of ocean sound level metrics using freeware software toolkits and developed core tools and processes that support open science. Four examples of comparative analyses that connect disparate PAM monitoring efforts, and integrate non-acoustic data illustrate how comparable, interoperable sound level metrics support a more coherent and synoptic perspective on global ocean soundscapes using methods that current and future PAM projects can leverage. Such a framework around PAM big data offers the opportunity to revolutionize large-scale marine ecology and oceanography in similar ways to other transformative approaches for understanding environmental or ecological patterns and processes at global scales.

## Introduction

Big data, characterized by its high volume, velocity, veracity, variety, and value, have revolutionized how we collect, analyze, and interpret information across diverse fields^1,2^. Passive acoustic monitoring (PAM), an example of big data, provides unprecedented opportunities to study soundscapes and their ecological and anthropogenic drivers across terrestrial, freshwater, and marine environments^3–7^. Specifically, marine PAM is employed throughout the global ocean by academic researchers, resource managers, industry, and military to understand how marine animals use sound^8–19^ and the impacts of anthropogenic noise on species and ecosystems^20–27^, including how ocean soundscapes are affected by change^7,28–32^ (**Table 1**). These diverse drivers for data collection coupled with cheaper and thus more accessible instrumentation, longer deployment periods and an increasing variety of deployed sensors have led to an exponential growth of PAM data over the past decade^33–35^. Progress towards building robust frameworks for curation, management, and dissemination of PAM are emerging to match the scale of data collection^36,37^. However, scalable methods for interpretation and comparison lag, hindering delivery of actionable metrics to meet diverse applications and this exposes a need to strategically expand collaborative efforts.

**Table 1.** Diverse applications of PAM, including examples of the user-community and their applications of the data.

| User Community | Applications for Passive Acoustic Monitoring Big Data | Example References |
| --- | --- | --- |
| Offshore industries | Offshore energy companies: siting, permit applications, monitoring plan designs, protected species compliance<br>Ports and maritime transport organizations: routing, traffic assessments | ZoBell, Vanessa M., et al. "Retrofit-induced changes in the radiated noise and monopole source levels of container ships." *Plos one* 18.3 (2023): e0282677. |
| Federal action agencies | BOEM: NEPA environmental assessment for offshore development<br>U.S. Navy and U.S. Coast Guard: Testing/training and port access | Van Parijs, Sofie M., et al. "NOAA and BOEM minimum recommendations for use of passive acoustic listening systems in offshore wind energy development monitoring and mitigation programs." *Frontiers in Marine Science* 8 (2021): 760840. |
| State and federal natural resource agencies | NOAA Fisheries: Integrated Ecosystem Assessment, essential fish habitat/habitat of particular concern-spawning fish monitoring, Endangered Species Act/Marine Mammal Protection Act-cetacean monitoring<br>NOAA ONMS: condition reports, management plans; California Ocean Protection Council, West Coast Regional Alliance<br>New York State Department of Environmental Conservation: New York Bight Whale Monitoring Program | Bolgan, Marta, et al. "Use of passive acoustic monitoring to fill knowledge gaps of fish global conservation status." *Aquatic Conservation: Marine and Freshwater Ecosystems* 33.12 (2023): 1580-1589. |
| Academic and industry big data users | Machine learning and artificial intelligence techniques applied to oceanographic and ecological studies | Frasier, Kaitlin E. "A machine learning pipeline for classification of cetacean echolocation clicks in large underwater acoustic datasets." *PLoS Computational Biology* 17.12 (2021): e1009613. |
| Cross-federal to international organizations | Council on Environmental Quality-SOST Interagency Ocean Sound and Marine Life Task Force COVID soundscape study<br>United Nations International Maritime Organization's ship quieting initiatives<br>International Quiet Ocean Experiment Science Plan initiatives | Miksis-Olds, Jennifer L., et al. "Envisioning a global multi-purpose Ocean acoustic network." *Marine Technology Society Journal* 55.3 (2021): 78-79. |
|  | Global Ocean Observing System (GOOS) Essential Ocean Variable implementation plan |  |
| Academic acousticians | Project PIs who collect PAM data for research purposes and/or teaching students, or for science communication | Parsons, Miles JG, et al. "A Global Library of Underwater Biological Sounds (GLUBS): An Online Platform with Multiple Passive Acoustic Monitoring Applications." *The Effects of Noise on Aquatic Life: Principles and Practical Considerations*. Cham: Springer International Publishing, 2024. 2149-2173. |
| Environmental consultants | Companies who provide services that could encompass characterizing industrial noise, biological sounds, and shipping activity | Austin, Melanie E., S. Bruce Martin, and Craig R. McPherson. "Measurements of Underwater Radiated Noise from Mobile Offshore Drilling Units." *The Effects of Noise on Aquatic Life: Principles and Practical Considerations*. Cham: Springer International Publishing, 2024. 1683-1696. |

With this large variety of PAM users and uses (**Table 1**), it is not surprising that tremendous progress has been made by individual projects, spanning regional to national scales and across multiple years, to develop practices for data curation, management, integration, and visualization. While not exhaustive, **Box 1** provides examples of the progress made in these areas. As the PAM community moves toward international standardization for soundscape measurements, several best practices documents were published^38–40^ and have been implemented among long-term monitoring projects (e.g., Miksis-Olds et al.^41^). To avoid duplication and divergent products, we seek to promote the use of centralized assets (i.e., data and tools) where appropriate, leverage what has already been built for a larger group of stakeholders, and coordinate further development opportunities.

#### Box 1.

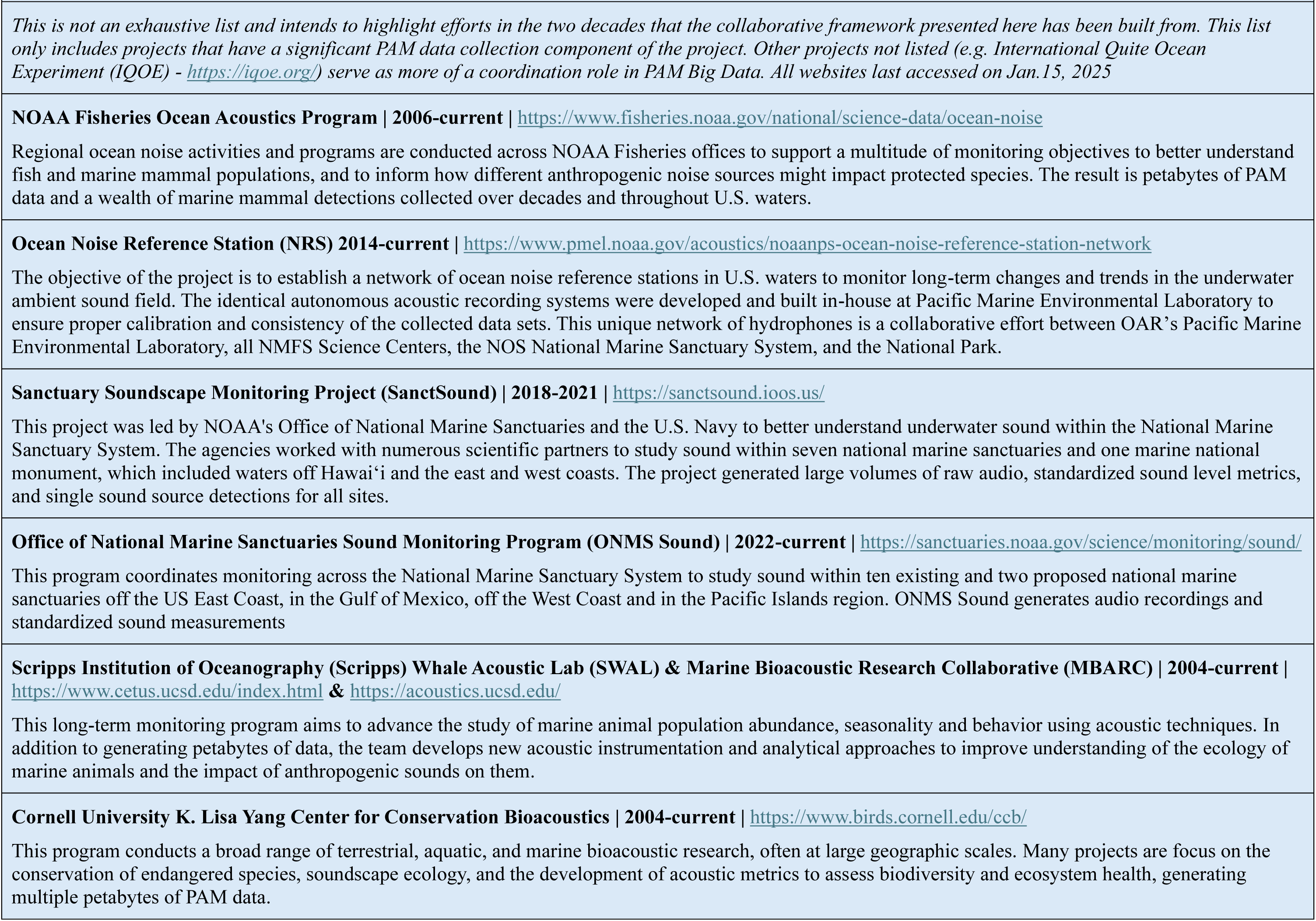
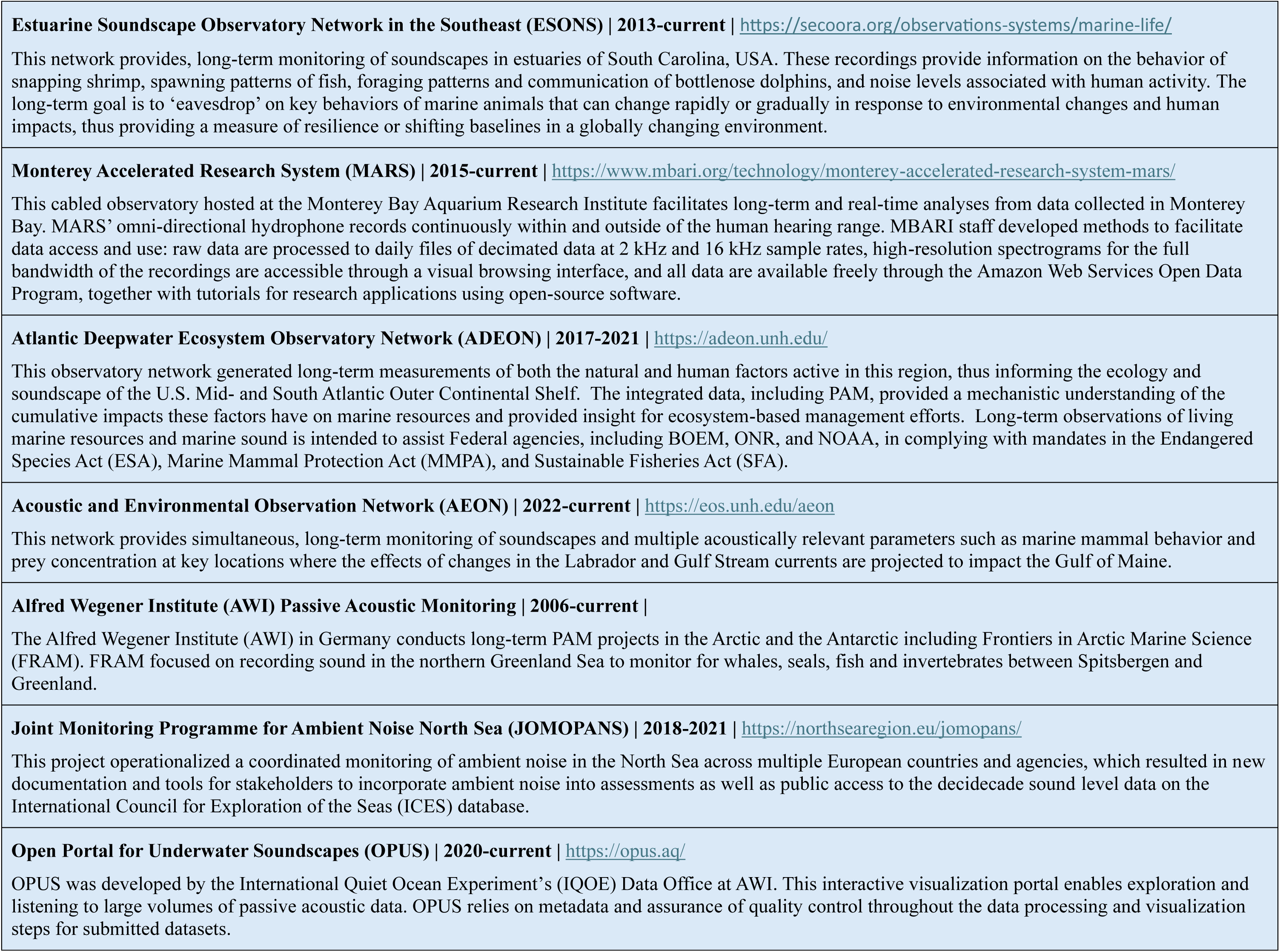
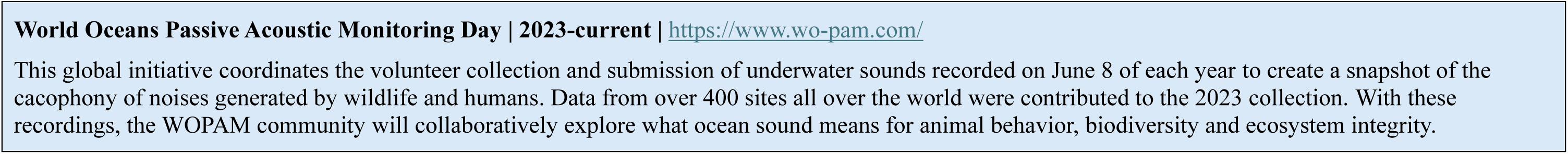
Passive Acoustic Monitoring Big Data Projects.

The Sound Cooperative (*SoundCoop*) project was created to address challenges in PAM curation, processing, and sharing by developing innovative, community-oriented tools and processes that make it easier and more efficient to extract information from large datasets and compare across monitoring programs. Funded by three U.S. federal government agencies (i.e., National Oceanic and Atmospheric Administration [NOAA], Bureau for Ocean Energy Management [BOEM] and U.S. Navy), *SoundCoop* leveraged existing national and international datasets and management efforts, identified remaining gaps, and built capacity towards community-prioritized functionality. The project’s framework focused on cyberinfrastructure that 1) created comparable, standardized sound level metrics using freeware software for a variety of recording equipment; 2) developed a standardized file format for an interoperable metric output; 3) established workflows to access data from separate cloud repositories; 4) co-visualized sound levels and integrated them with environmental data in a performant, interactive public portal; and 5) published open-source tools for the community to replicate the project’s processing and visualization methods. The success of such applications relies on the ability to develop standard and broadly accessible data management mechanisms, and to continue to ensure that these mechanisms can scale effectively with increasing global participation. Here, we showcase the progress achieved through *SoundCoop* in addressing big data challenges for PAM, presenting the project as a model framework for tackling similar challenges across scientific fields.

## Building a Collaborative Framework

The *SoundCoop* project’s collaborative approach provided a unique opportunity to address the challenges associated with big data in marine PAM by focusing on existing community standards, knowledge, and datasets and thus established a community-driven process. Previous efforts with PAM demonstrated the value of exploring datasets from separate monitoring projects by leveraging their centralized availability, and processing the data into a standardized and comparable metric^42^; however, the workflow was built solely to meet the needs of that project-specific scientific objective, and was not necessarily generalizable for broader meta-analysis or community adoption. By engaging a broad network of stakeholders, *SoundCoop* fostered collective input to identify key obstacles, and develop practical and scalable solutions. The project emphasized the creation of standard outputs from freeware tools, ensuring consistency and accessibility for numerous PAM user groups. Additionally, it championed the use of public repositories and data portals to promote transparency, data sharing, and reproducibility. This framework serves as an example for harnessing collective expertise to identify a meaningful data product to address the heterogeneity in raw data and enable comparison across monitoring projects; efficiently share and connect data to address the challenges of dissemination associated with large data volumes; integrate data quality information to address noise accumulation; and streamline access to federated datasets, especially cloud-based to enable further processing capabilities in scalable, high compute environments.

### Broad Community Input

Given the international scope, multiple U.S. federal agency applications, and diverse applications (**Table 1**), the *SoundCoop* effort was guided by an advisory committee and several stakeholder engagement events. The advisory committee represented experts across federal agencies, academic institutions, and industry, namely the U.S. Navy’s Living Marine Resources Program and Office of Naval Research, BOEM, NOAA National Marine Fisheries Service (NMFS), NOAA Office of National Marine Sanctuaries, NOAA Integrated Ocean Observing System (IOOS), IOOS’ Northeastern Regional Association of Coastal Ocean Observing System, University of Colorado Boulder / NOAA NCEI, Axiom Data Science, Monterey Bay Aquarium Research Institute, the Regional Wildlife Science Collaborative for Offshore Wind, Alfred Wegener Institute in Germany, International Quiet Ocean Experiment, and Rijkswaterstaat Ministry of Infrastructure and Water Management in Netherlands.

During the first phase of community input, the project’s steering committee collated and reviewed information on the status of existing national and international PAM data management tools in support of *SoundCoop*’s focus on enhancing longer-term and large-scale curation, access and operability. Previously collected PAM datasets were selected by the team that represent 12 disparate monitoring efforts with varying collection purposes and applications, 10 recording instruments deployed across 7 geographic regions spanning 17 year and varying sampling regimens. Datasets were grouped into four categories:

1. Long-time series recorded in the Arctic. Data were contributed from the NOAA NMFS Alaska Fisheries Science Center Arctic Long-Term Integrated Mooring Array (AFSC-ALTIMA), NOAA-National Park Service Ocean Noise Reference Station Network (NRS), and U.S. Navy-funded Scripps Whale Acoustic Laboratory (SWAL) projects^43–45^.
2. Variation in recordings throughout 2021 across three geographic regions: Gulf of Maine, Southeast US, and Central California Coast. Gulf of Maine datasets were contributed from the Acoustic and Environmental Observation Network (AEON) program, NOAA-Navy Sanctuary Soundscape Monitoring (SanctSound) project, and NOAA NMFS Northeast Fisheries Science Center (NEFSC) monitoring project^46–48^. The Southeast US dataset was recorded in the May River from the Estuary Soundscape Observatory Network in the Southeast (ESONS) project^49–51^. The Central California Coast datasets were from MBARI Monterey Accelerated Research System (MARS) and the NRS project^44,52^.
3. Offshore wind monitoring. Sound data were contributed from a site off Virginia (Mid Atlantic) that was part of a BOEM-funded, Cornell University-led project as well as North Atlantic right whale (*Eubalaena glacialis*) upcalls detected off the New York Bight from a New York State Department of Environmental Conservation-funded, Cornell University-led (NYSDEC-Cornell) project^53–55^.
4. International comparisons. Two international (non-U.S.) datasets were contributed from AWI’s Frontiers in Arctic Marine Monitoring (FRAM) and the multiple European Union agency-led Joint Monitoring of Ambient Noise in the North Sea (JOMOPANS) projects^56–60^.

An in-person workshop held a year into the project served as a key opportunity to check in with *SoundCoop* participants. Since many who shared data were not directly involved in the mechanics of the project, it was important to be transparent about progress towards the project goals while also allowing participants to access the tools being developed and see data product visualizations. Through guided discussion and hands-on activities, participants were able to share feedback to the steering committee.

During the third phase of community input, a hybrid-format workshop was held to allow broader participation especially from those working more broadly on big data challenges (e.g., *NCEI World Ocean Database, NOAA Center for Artificial Intelligence, IOOS-funded Reaching for the Cloud project*). The first session summarized the successes of this large undertaking, highlighting the enormous community efforts as well as key advances in metadata and data quality, and robust and repeatable methods. Sharing with a broader big data community raised awareness of complementary efforts, and helped ensure longevity and scalability of the progress made in *SoundCoop*. The final session focused on future pathways and included use and application demonstrations from different user communities (**Table 1**). These groups are currently building upon what was established in *SoundCoop* to operationalize the creation of standardized sound level metrics in an interoperable self-describing format for their programs. Lastly, we see this collaborative publication as the final phase to broad community input by creating a collective vision we can all build from.

### Standard Outputs from Open-Source Tools

Standard outputs from open-source tools benefit big data methodologies by ensuring consistency, accessibility, and efficiency in data analysis, and promote collaboration and sharing^1^. Leveraging existing documentation and associated software for a broader community of PAM practitioners, all *SoundCoop* data (except for the NYSDEC-Cornell and JOMOPANS datasets) were processed into daily files of one-minute resolution hybrid millidecade (HMD) spectra^61,62^. This method balances reduced data volume with adequate resolution of acoustic spectra^61,62^. This soundscape metric was used here as a way to provide a high level, comparison-ready overview of data that are too large to explore directly. Further, open-source and freeware processing tools were chosen to calculate the HMD spectra to ensure the broader community can replicate the work without the barrier of paid licensing. The *Making Ambient Noise Trends Accessible* (MANTA) Matlab tool and the *Python passive acoustic analysis tool for Passive Acoustic Monitoring* (PyPAM) with PyPAM-Based Processing wrapper (PBP) were both used for processing^63–65^. Collectively, over 14 million minutes of recordings were processed into HMD spectra, equating to nearly 30 years of marine sound levels. Participants preferred multiple software options to provide flexibility for the community with diverse processing capabilities and technical expertise. As a result, careful evaluation throughout multiple phases of the processing, from calibration to content of the output were required. Summaries of the processing methods and decisions are found in **Appendix I**.

To ensure interoperability in the software output and facilitate data sharing, a standardized netCDF file with complete, self-describing metadata was developed. Content on the netCDF structure is found in **Appendix I** and metadata details are described in **Appendix II**. This new standardized output for PAM sound level metrics provides a common, standards-driven format that simplifies integration across diverse datasets, reducing the time and effort required to process and analyze data from multiple sources. Included within is an effort variable that documents the number of seconds of acoustic recording that went into each HMD minute. The netCDF also includes a digital object identifier (DOI) for the respective project provided by the NOAA NCEI Passive Acoustic Data Archive, where the datasets are stewarded. DOIs are persistent identifiers that facilitate transparency, long-term stewardship, and citability of the datasets. Workflows built using open-source tools, which are generally widely accessible, that input these standard, machine-readable netCDFs further lower barriers for different user groups^66–68^. Moreover, standardized outputs facilitate transparency and reproducibility, critical for fostering collaboration in big data projects^69,70^.

### Public Repositories and Data Portals

Public repositories and data portals provide critical infrastructure for maximizing the utility of big data by ensuring accessibility and facilitating open science^71,72^. By hosting globally accessible data in standardized formats with complete, standards-driven metadata, data repositories enhance data discoverability and interoperability. This enables integration with other publicly accessible datasets and open-source tools, and ultimately meets FAIR (Findable-Accessible-Interoperable-Reproducible) principles^73,74^. The comparable HMD spectra produced through *SoundCoop* were distributed across three freely accessible platforms: NCEI Passive Acoustic Data Archive Google Cloud Platform (GCP) bucket (<u>link</u>), MBARI Amazon Web Services (AWS) S3 bucket (<u>link</u>), and Axiom Data Science Research Workspace (<u>link</u>). Automated workflows leveraging Python libraries were developed to programmatically access files from the GCP, AWS, and the Axiom’s Workspace, and published as a community resource (<u>link</u>). Ultimately, these repositories democratize data access, empowering a broader range of stakeholders—including policymakers, educators, and community organizations—to leverage big data for informed decision-making and innovative solutions.

To promote discoverability and demonstrate the value of standardized products for PAM housed in public repositories, an interactive data portal was developed that visualizes the *SoundCoop* datasets and integrates non-acoustic data from marine environmental sensors (<u>link</u>). Specifically, wind speed and wave height were accessed from the IOOS environmental sensor map (<u>link</u>) and integrated into the *SoundCoop* data portal. The one-minute HMD spectra were divided based on categories of wind speed and wave height that align with sea state: wind speed categories were < 10 kts (Beaufort scale 0-3), 10 – 20 kts (Beaufort scale 4-5), and greater than 20 kts (Beaufort scale 6+) and significant wave height categories were <1 m (Beaufort scale 0-3), 1-3 m (Beaufort scale 4-5), and greater than 3 m (Beaufort scale 6+). One-degree resolution sea surface temperature values for a given site and month were extracted from the NOAA World Ocean Atlas (WOA) 2023^75,76^ to compute sea surface temperature climatologies for each site. Sea surface temperature anomaly was then calculated by subtracting the WOA 2023 derived temperature climatology from each measured sea water temperature value at that site for the recorded month. The integration of publicly available sensor data provided a first-tier interpretation of the sound level data, leading to some initial insights into patterns within the acoustic data. These data portals provide tangible examples of the benefit of standardized data products for big data projects, like PAM, by enabling novice users to explore large volumes of data with ease, fostering curiosity, and supporting learning while also empowering experts to identify areas for deeper analysis.

## Value of Comparative Analyses

The value of big data lies in its potential to transform raw values into actionable insights, identify emergent properties that can drive decision-making and innovation^72,77,78^. In today’s data-driven world, extracting value from big data enables organizations to uncover patterns, predict trends, and advance management objectives across diverse applications^1^. In the context of PAM, this includes ensuring species protection^35^, reducing noise in protected places^79^, and understanding the blue economy activity in coastal and offshore ecosystems^80^. However, realizing this value depends on effective data management, advanced analytical tools, and a clear understanding of how insights align with organizational goals or societal needs. By focusing on value, big data becomes more than just vast quantities of information—it becomes a powerful catalyst for informed decisions and progress.

Within this paper, we incorporated a subset of the overall *SoundCoop* datasets to provide concise examples of the versatility of HMD spectra and the benefit of comparable sound level metrics (**Appendix III**). Case Study 1 examines changing sound levels relative to a reference period. Case Study 2 compares sound levels at regional to global scales. Case Study 3 discovers dominant sound sources through indices. Finally, Case Study 4 advances interpretation of the sound levels through integration of environmental data. Collectively, these comparisons elucidate important components of our ocean soundscapes, how they vary by region and over time, and the utility of the HMD data product for insights from PAM data. All analyses were performed in Python using Jupyter Notebooks run on a local JupyterLab. The Jupyter Notebooks can be found in the SoundCoop project’s Github repository (<u>link</u>) and visualizations of the datasets can be found in *SoundCoop* data portal. The location of all SoundCoop sites is illustrated in **Figure 1**.

**Figure 1.**
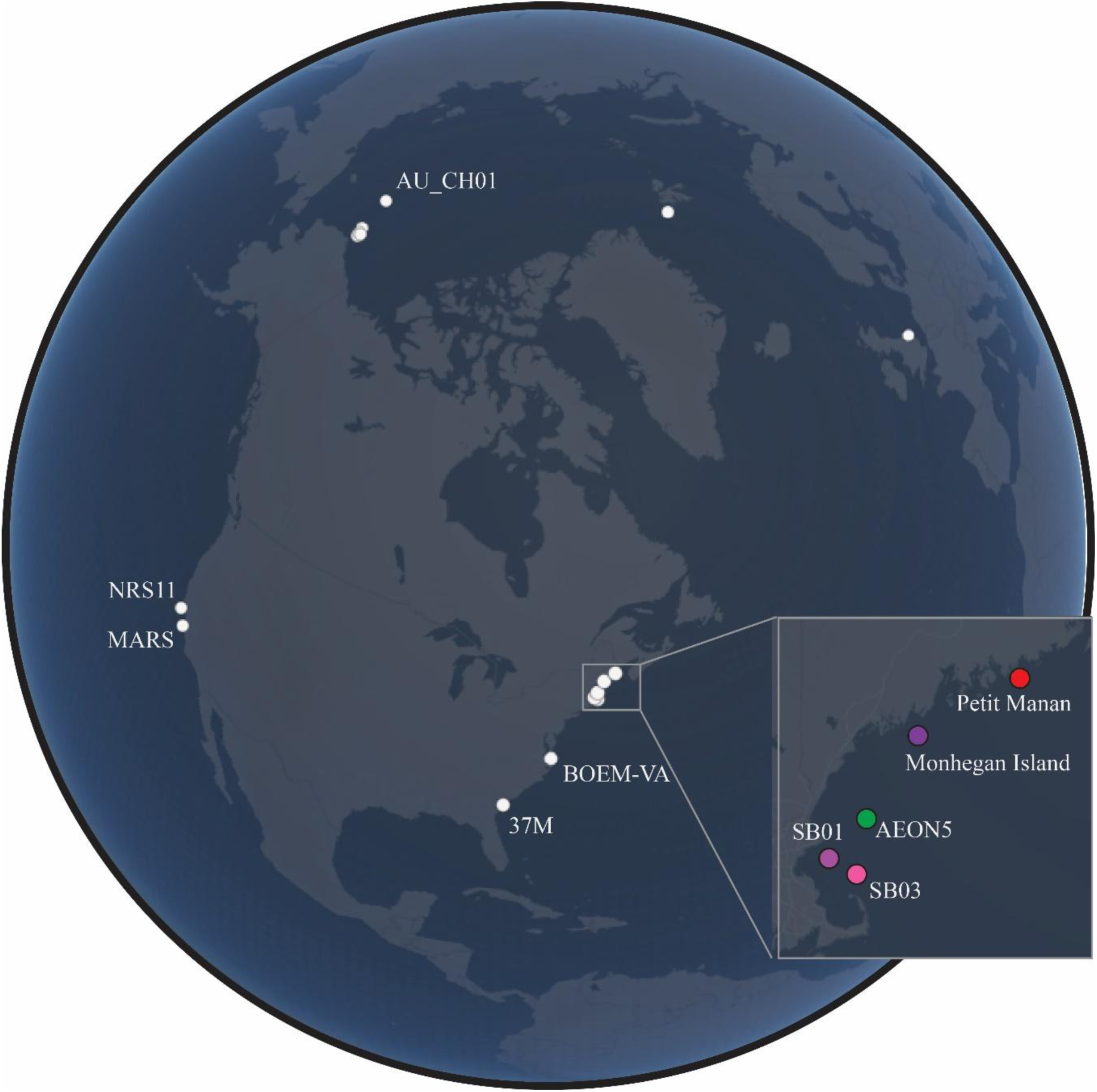
Location of all SoundCoop recording sites. Sites that are labeled are included in the Case Study analysis presented in this paper. The colored dots for the sites in the Gulf of Maine inset correspond to the colored lines in the power spectrum plots in Figures 3 and 4.

### Case Study 1: Change Relative to Reference Period

When a PAM program has been maintained over a long period of time, the resulting time series can be used to show how sound levels may be changing in a specific location on seasonal, annual, and interannual scales^81–85^. The AFSC-ALTIMA project has recorded sound in the Arctic since 2007 to monitor marine mammal distribution and migration timing^86^. The project’s AU_CH01 site is used here to compare how sound levels between 100 Hz and 1,000 Hz from HMD spectra vary over time (**FIG. 2)**. To control for the influence of seasonal variation, including the influence of sea ice coverage on the soundscape, only sound levels from September months, a constant ice-free period at this location, were included.

**Figure 2.**
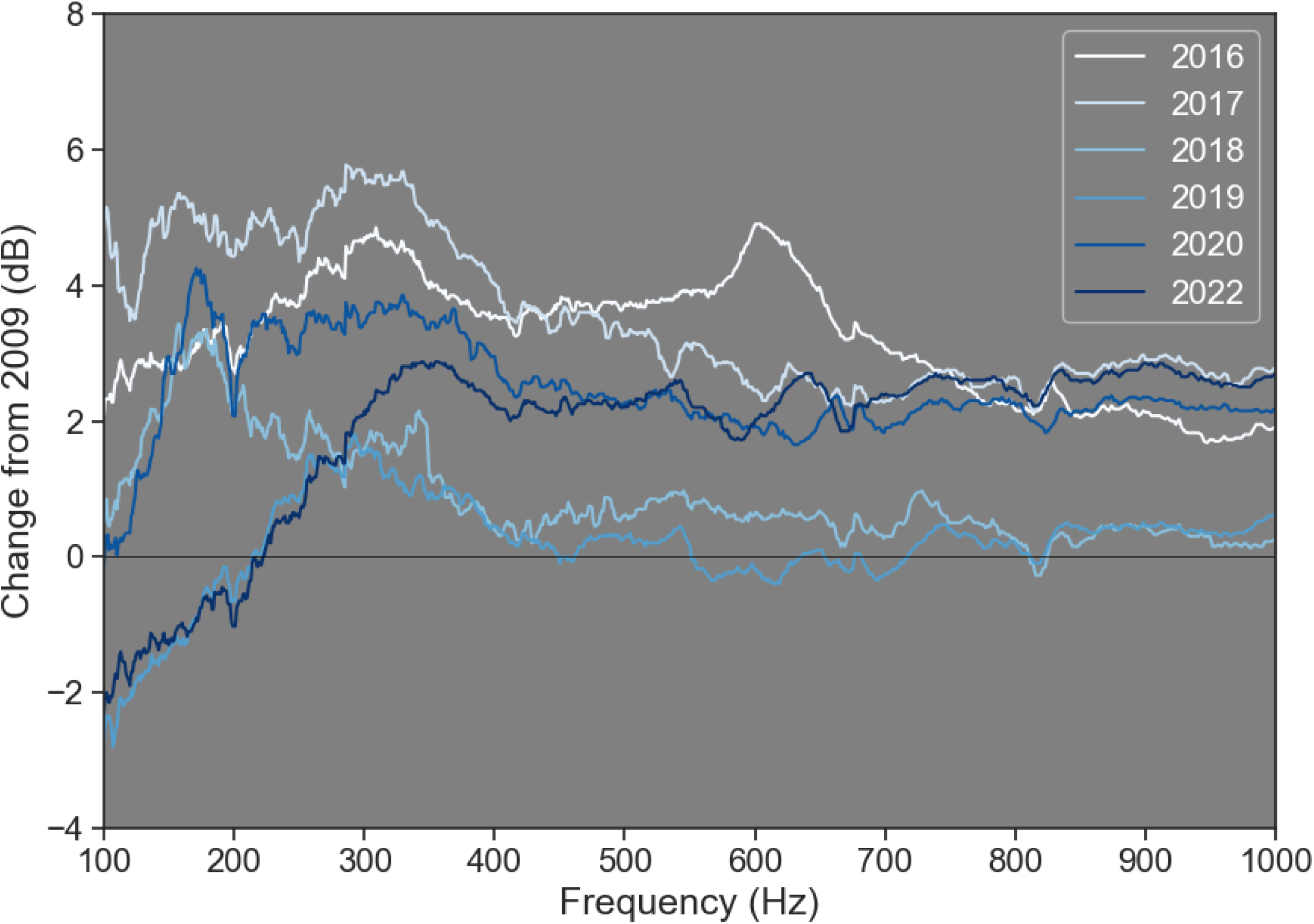
Change relative to reference period at a long-term PAM monitoring location in Arctic Ocean at site AU_CH01. Change in the month of September, an ice-free period, is shown, and a reference year was selected, defined here as the earliest year in which there were over 80% “Good” quality minutes available, specifically 2009. The one-minute HMD spectra were median aggregated to one-day resolution, and processed into a median power spectral density (PSD) for each available year’s September. A 5-kernel window size, one dimensional median filter was applied to the median PSD to remove impulsive artifacts. Years with “Good” quality minutes less than 80% of the expected total were omitted from the analysis. This threshold was applied to 2008, 2010-2015, and 2021. The September 2009 median PSD was subtracted from subsequent year’s September median PSD, and plotted to illustrate relative change across years. The frequency range is truncated to 100 to 1,000 Hz.

The comparison presented here provides initial insight on the dominant patterns of change in ocean sound. Generally, sound levels recorded above 300 Hz were higher in amplitude for years following 2009 with 2016 and 2018 reaching over 4 dB higher. Lower frequency (100-300 Hz) sound levels were up to 2 dB lower in 2019 and 2022 compared to 2009 and up to 7 dB lower than 2018. Sounds from biological (e.g., migrating whales), physical (e.g., higher sea states due to storms), and anthropogenic (e.g., noise from vessels) sources all contribute to the overall soundscape, therefore, these broad patterns in sound levels over 7 years inform additional analysis, including what frequency bands to focus on.

### Case Study 2: Comparisons at Regional to Global Scales

A major benefit of producing a robust workflow to create standardized sound level metrics in an interoperable format is that datasets from different monitoring programs across federated repositories are now easily comparable. To illustrate this concept, we co-visualized HMD spectra from eight sites (AEON5, 37M, MARS, Petit Manan, Monhegan Island, NRS11, SB01 and SB03) recorded across three geographic regions (Gulf of Maine, Southeast U.S., and Central California Coast). Data were pulled from three different data repositories (GCP, AWS, and Axiom Workspace), processed into weekly-aggregated median PSD that represent sound levels recorded throughout 2021, and visualized (**FIG. 3**).

**Figure 3.**
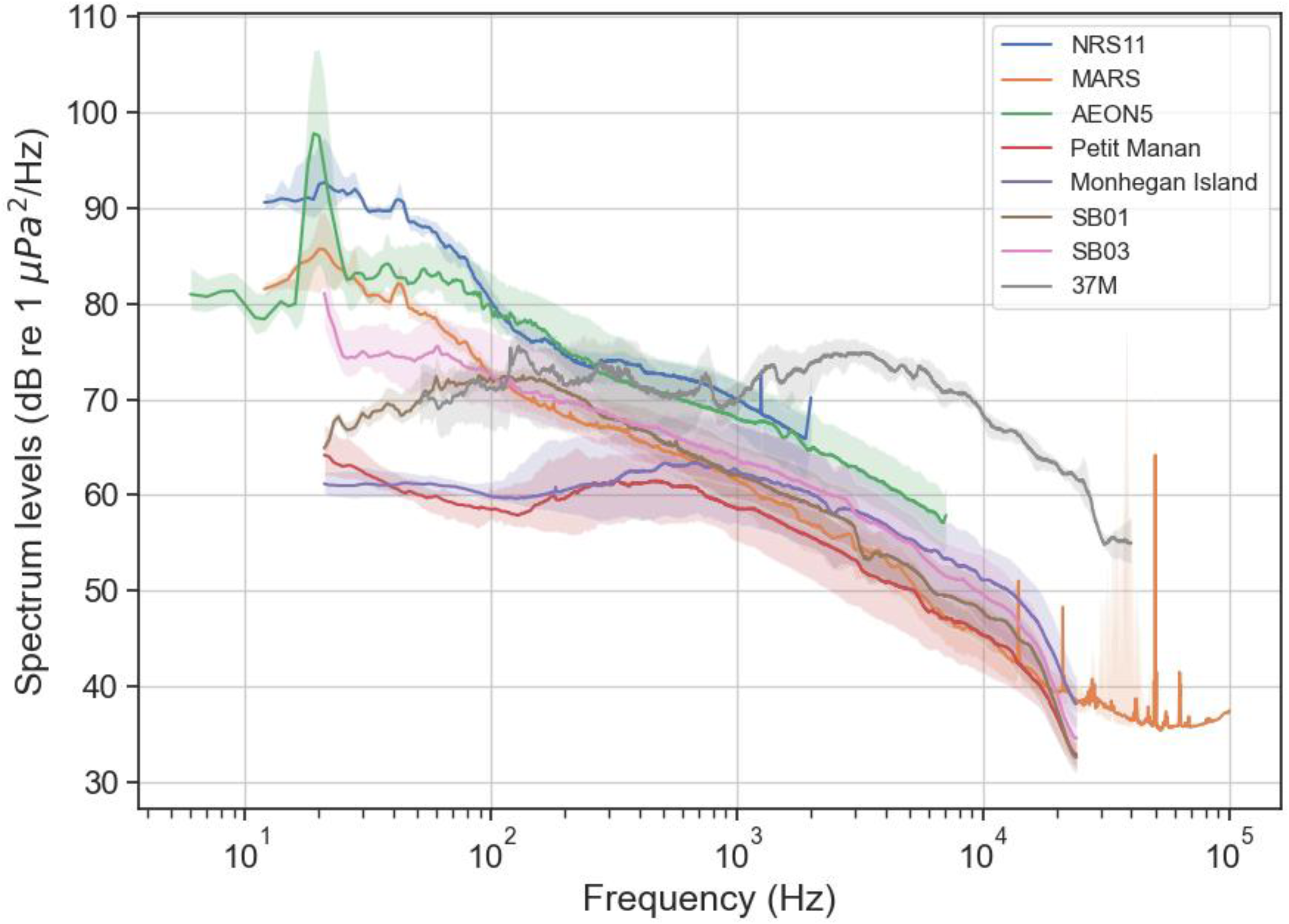
Large-scale comparisons of weekly-aggregated median power spectral density for eight sites recording throughout 2021. Sound levels are shown as the median (solid line) with 25th and 75th percentiles (shaded area).

While the value demonstrated here mainly focuses on the interoperability of standard data products, broad differences emerge from this comparison, relating to the oceanographic settings. In the Gulf of Maine, sound levels at the deepwater AEON5 site were consistently higher than the shallower SB01 and SB03 sites, likely due to the more efficient sound propagation in deeper water and the sites’ close proximity to a heavily-used shipping channel with traffic from both large and small vessels^21,25^. Petit Manan and Monhegan Island sites had the lowest overall sound levels likely attributed to their coastal location away from the shipping channel and large vessel traffic. Located at the mouth of May River where there is heavy small vessel traffic and significant fish sound production^50,51,87–92^, ESONS’ 37M sound levels were high between 100-1000 Hz mostly due to snapping shrimp snaps, fish sound production, and vessel noise, and highest across all sites above 1000 Hz due to snapping shrimp snaps as well as vessel noise and silver perch chorusing^51,90,92^. The two Central California Coast sites, NRS11 in the Cordell Bank National Marine Sanctuary and MARS in the Monterey Bay National Marine Sanctuary, show elevated sound levels at lower frequencies, which are mainly attributed to the presence of larger vessels and seasonal migration of whales^93,94^.

### Case Study 3: Dominant Features Revealed

Producing standardized data products provides opportunities to target additional analyses that leverage the original quality-controlled product. The result is further insights that can be compared across space and time, although considering context for the analysis becomes even more relevant as further indices are derived^95^. To highlight what sources could be contributing to the soundscape and how their presence varies throughout the year at different locations, we leveraged established methods to identify blue whale (*Balaenoptera musculus*) B calls and fin whale (*B. physalus*) 20 Hz calls for deepwater recordings^14,93,96–99^. These methods use a call index (CI) to quantify the intensity of the calls relative to background noise applied here using the minute-resolution HMD netCDFs from the *SoundCoop*’s deepwater sites (> 100 m): AEON5, MARS, and NRS11.

The blue and fin whale call indexes show seasonal patterns across the sites (**FIG. 4**). For example, blue whale B calls, as determined from the call index, are present in the NRS11 and MARS recordings in winter (January to March) and late summer to winter (August through December). MARS has two peaks later in the year in August and again in November while NRS11 only has a single peak around October. Fin whale calls as identified in the call index were detected across all three sites with the strongest peaks occurring in the fall. These patterns align with previous findings for blue and fin whale sound production^11,14,93,97^ and highlight the versatility of the HMD spectra to be adopted for existing algorithms due to its high temporal resolution and resolution at low frequencies.

**Figure 4.**
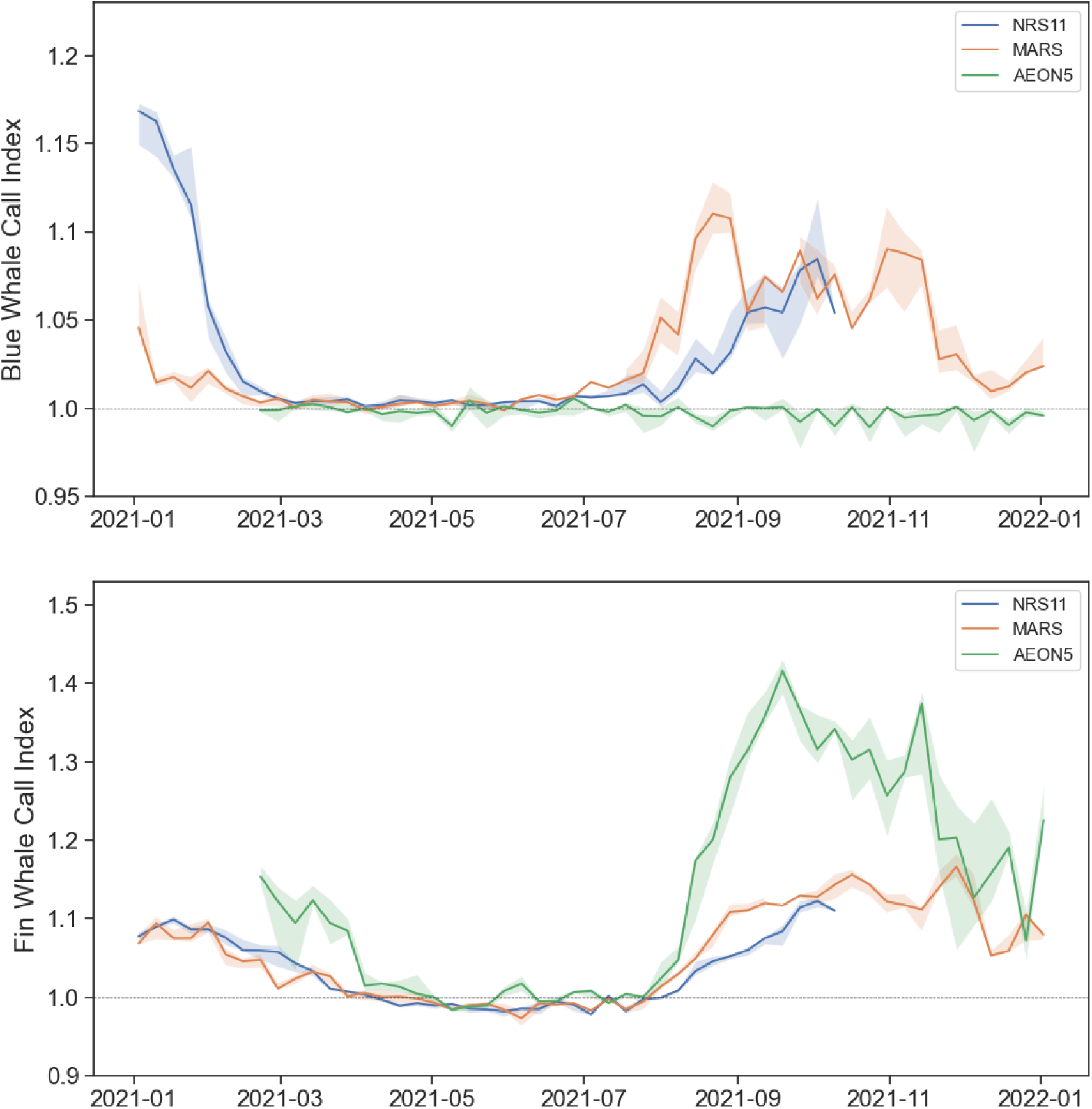
Comparison of blue and fin whale call indices derived from standard products. Weekly whale call index for (top) whale B calls and (bottom) fin whale 20 Hz calls for three deeper water sites. The solid line shows the median (50th percentile) CI value for that week with the shaded envelope delineating the 25th and 75th percentiles. The dotted line at 1.0 denotes where the signal would be above background noise (above the line) and where the signal would be less than the background noise (below the line). For blue whales, the CI ratio was calculated as [42, 43 Hz] / [37, 50 Hz] where the 3rd and strongest harmonic of B calls occurs between 42 and 44 Hz, and 37 and 50 Hz represent frequency bands that would not include energy from a B call. The results are then binned to one-day resolution using the median. CI was calculated for fin whale calls using [20, 21 Hz] / [12, 34 Hz]. The CI results were then aggregated further to one week resolution and plotted over time.

### Case Study 4: Integration with Environmental Data Sources

While PAM is still emerging in the big data landscape, monitoring of the physical environment at global scales is well established (e.g. wind speed, wave height)^100–104^. These physical conditions provide insight on drivers of changes in soundscape. For example, surface agitation from wind and waves add acoustic energy to the ocean with known effects on the sound levels^105^. To demonstrate the value of data integration, HMD data at a PAM site in the North Atlantic near the mouth of Chesapeake Bay (BOEM-VA) was combined with wind speed, wave height, and sea surface temperature pulled from an open-source data repository (‘gov-ndbc-44014’ ERDDAP dataset). During a two-week period (Aug 28 - Sep 20, 2016), the integration revealed the influence of Hurricane Hermine^106^ on the soundscape (**FIG. 5**). Specifically, sound levels at this site were highest when Beaufort Sea State was 6+, which is associated with wind speeds greater than 20 knots and wave height greater than 3 m. Sound levels were most strongly influenced by increased wind speed and wave height at lower frequencies (less than 50 Hz) and frequencies above approximately 800 Hz. A decrease in noise from vessels occurs prior to the storm as a result of Tropical Storm Warnings^107^, and changes in biological sounds also emerge. During the storm, diurnal fish chorusing events between 100 and 1,000 Hz dramatically increase in amplitude and an additional chorusing event (likely a different species) between 1,500-2,700 Hz begins. Beyond this specific event, continuous data integration of the physical environment with sound levels provides key insights on the seasonality of soundscapes dominated by physical conditions^95^ and could help monitor the interplay of ocean sounds and wind-wave dynamics in a changing climate^108,109^. As other sources of big data become available, scalable integration with PAM data becomes possible furthering the ability to understand drivers of change in ocean soundscapes.

## A Collaborative Way Forward

Through this collaborative multi-year project, big data challenges in PAM were overcome and a framework to progress the community’s collective analytical capabilities emerged. The project built on previous efforts (**Box 1**), brought together a broad community of practitioners (**Table 1**), established scalable processing tools and pathways^63,64^, and demonstrated the value of comparisons (Figs 2-5). Key accomplishments include 1) creating comparable, standardized sound level metrics using open-source software that supports a variety of datasets, 2) developing a standardized, machine-readable sound level metric file format that ensures consistent file structure with embedded metadata for interoperability and dissemination, and 3) programmable access to free public repositories of sound level metrics and environmental sensor data through collaborative processing tools.

**Figure 5.**
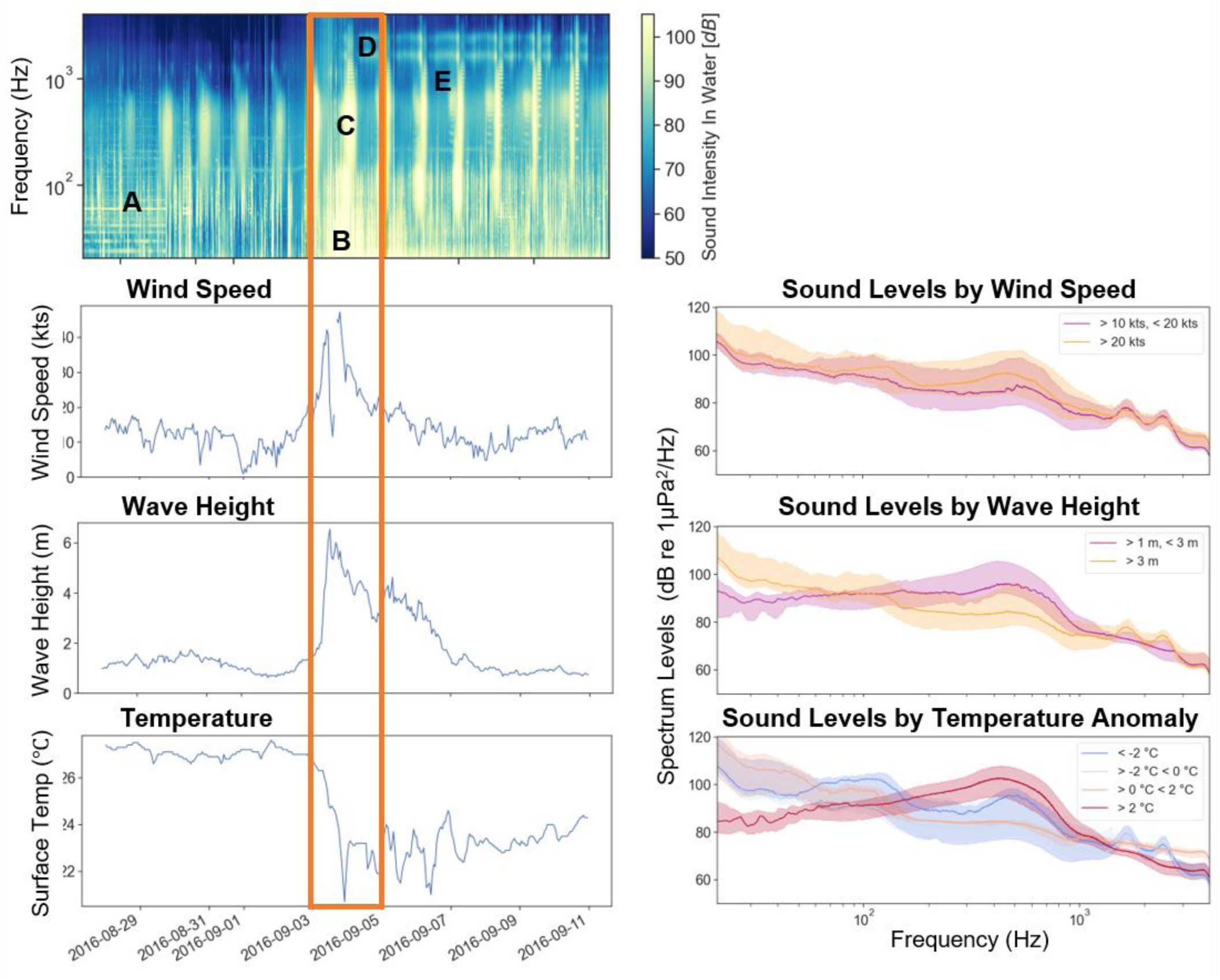
Integrated data sources reveal weather related changes in underwater soundscapes. The HMD spectra were median aggregated to one-hour resolution to match the hourly resolution of the wind speed, wave height and water temperature data (left panel) for the 2-week time period before, during, and after Hurricane Hermine was near the recording instrument, namely August 28 - September 10, 2016. Hourly median PSD were calculated for time periods associated with Beaufort scale 0-3, Beaufort scale 4-5, and Beaufort scale 6+, and temperature anomaly bins of < -2℃; -2℃ < 0℃; 0℃ < 2℃; > 2℃ when the storm was present in the area (September 3-4, 2016, which is highlighted in the orange box in the left panel and results plotted in the right panel) based on the NOAA National Hurricane Center hurricane/tropical storm track and the nearby NOAA National Data Buoy Center (NDBC) buoy (gov-ndbc-44014) wind speed. This buoy is located 82.9 km from BOEM-VA PAM site. Annotations in the spectrogram denote A) noise from vessel activity; B) low frequency noise from wind associated with the storm; C) nocturnal fish chorus of increasing intensity as storm nears and passes; D) new fish chorusing that begins after the storm passes, and E) third fish chorusing event that arose after the storm.

### Challenges Overcome

The sheer size of data produced by modern monitoring systems, like PAM, is vast, especially when monitoring over extended periods and across multiple locations^42,110^. *SoundCoop*’s new infrastructure enables transformative, translational research by improving the accessibility of data, information, and methods for standardized metrics and formatting. Free software toolkits facilitated the standardized processing of sound levels. However, extensive testing and revisions were needed to produce the necessary output of predictable and therefore comparable 60-second time bins, ensure accurate second counts within each bin, and handle the variety of recorder types associated with the *SoundCoop*. This effort resulted in a new suite of datasets to test against including different instruments, calibration considerations, “wiggle” in the internal timekeeping, and unexpected recording gaps. Tracking each second of every recording is crucial to produce accurate and reproducible one-minute resolution HMD files. This provides fine-scale accuracy while correct calibration ensures statistical importance to the analytical results. Comparing multiple variables, namely HMD spectra, time, and effort, from the software outputs to a “ground truth” dataset - as was done in this project - is necessary to examine the accuracy of each time bin.

Over 14,000 netCDF files were produced from the HMD-processed datasets in this project. Managing these many files from different sources in a CSV format would have been brittle and required additional spreadsheets or databases containing each dataset’s metadata. The self-describing netCDF embeds all file-, deployment-, and project-level metadata within each file, facilitating efficient management, sharing, and interpretation. Aligning the outputs of both software to the same netCDF structure ensured interoperability, comparability of results, and seamless integration into higher-level processing workflows.

Tutorials empower practitioners to leverage PAM data more effectively, while visualization tools and standardized workflows simplify large-scale data analysis, reducing the need for individuals to “reinvent the wheel.” To ensure transparency and reproducibility of the methods and results, and in turn provide tools that could be easily adapted to existing and future workflows, Python-based Jupyter Notebooks were developed. These notebooks document how to access, read in, aggregate, and visualize the project’s thousands of files using open-source processing in reproducible environments. The notebooks are shared publicly on GitHub along with documentation such as readmes and guidance for different user groups (from novice to expert, science-driven to management-driven). Environment setup files that capture system and library requirements were produced so that the notebooks could be run seamlessly by any user. Depending on the volume of data being processed and the processing needs, Google Colab, JupyterLab, Github Code Space, and Binder are potential options to run the notebooks for the community to explore – though there are assuredly many other environments that are best suited for individual needs including higher performing, cloud-based, fee-based platforms.

### Best Practices for Data Quality

Big data often includes unstructured, noisy, or incomplete data, making analysis complex^111^. To realize the value of PAM data, we highlight considerations when building big data frameworks to ensure data quality and interoperability between datasets. HMD offers a robust yet versatile foundational sound level metric product and these key requirements are necessary before applying this approach to your dataset:

#### Calibration

Proper characterization of the calibration is critical for accurate, absolute sound level measurements. Users must understand the calibration process and how the HMD software applies it. Clear documentation, intuitive workflows, and graphical outputs from the software are essential to vet this information. If calibration is unknown, not possible, or the hydrophone sensitivity degrades over the duration of the recording, processing the data into sound level metrics like HMD for quantitative analyses will not produce accurate results.

#### Timekeeping

Clock drift occurs in PAM recording units, and can accumulate over long deployments depending on internal clock quality and in response to temperature changes^112^. If drift is high, it can noticeably impact the accuracy of one-minute resolution HMD products if not accounted for. Autonomous recording systems are built to be fault-tolerant but errors can occur, leading to timekeeping problems such as lost or extra samples, or variability in true sampling rates. Each scenario needs to be properly handled by the software calculating the HMD, and a thorough examination of where each second is being accounted for in the calculation will ensure the one-minute time bins are accurate. In severe cases where timekeeping is highly variable, consider aggregating to a coarser time resolution (e.g., hour) or another metric that is robust to some temporal inaccuracy.

#### Data Integrity

Data quality must be thoroughly documented for proper use or reuse of sound level metrics. Within the frequency dimension, documentation should note where calibration is not available or where recording instrument accuracy limits the quantitative analysis of specific frequencies. This is particularly necessary for data processed with MANTA, which outputs 0 Hz to Nyquist regardless of the calibration range or presence of electrical noise. Within the time dimension, document periods of known issues and the timing of deployment and recovery. As demonstrated in *SoundCoop*, it is recommended to implement a machine-readable method for masking unverified, compromised, or poor-quality times and frequencies. This can be achieved by creating a numeric matrix of data quality tags that matches the dimensions of the power spectral density matrix. The data quality tags could derive from a standardized JSON metadata file, such as the one outputted by the NCEI PassivePacker PAM data documentation tool (<u>link</u>), or built with automated scripts following a quantitative QA/QC review of the processed data. The netCDF file with the HMD spectra results and data quality mask can then be used to programmatically ensure only “good” time periods and frequency bands are used in quantitative analyses.

These collective advancements foster collaboration and momentum within the PAM community, building partnerships and aligning efforts. By coordinating and standardizing data, the community can better address critical questions about the impacts of industrialization on marine habitats and life and enable scientists to explore climate effects on marine ecosystems in the most sustainable ways possible. This distributed model, where standardized tools are widely adopted, allows more researchers to contribute to collective solutions and deliver valuable insights to science and management applications.

### Emerging Big Data Considerations

The demand for real-time data analysis is increasing, especially in scenarios requiring immediate decision-making^113^. Real-time processing of acoustic data is particularly important in urgent scenarios such as detecting the changing presences of endangered species or responding to illegal activities like poaching or illegal fishing^114–117^. However, real-time analysis is computationally intensive and requires sophisticated algorithms and hardware to handle the continuous flow of data without delays. Notably, advances discussed here can directly support decisions that are neither immediate nor decadal but rather responsive to increasingly dynamic inter-annual changes in animal and human use patterns in ocean and coastal waters.

Managing access, privacy, and security concerns is critical when handling sensitive or proprietary big data^118^. Some PAM data may be sensitive, particularly in the context of national security, monitoring protected areas or species, and data sovereignty considerations arise with diverse data collection. Ensuring secure data transmission, storage, and sharing across platforms is important to protect both species and project confidentiality. Screening and filtering of some data may be necessary prior to public dissemination, but such treatments can be surgical and allow the majority of the collected data to inform a broad suite of users.

Providing public access to data, particularly data funded by government agencies, is acknowledged to increase transparency and citizen engagement, and to support economic growth through innovation, evidence-based policy making, and the ability for researchers and businesses to leverage data for new insights and solutions. However, providing public access to data will continue to raise ethical questions about data ownership, usage, and the potential consequences of insights derived from analysis^119^. In the context of PAM, although there are many conservation goals that can be advanced significantly through more efficient and interpretable data practices, there is also the potential for data misuse in ways that could harm protected species (e.g., revealing the location of fragile managed fish or mammal concentration). Considering such risks while empowering new applications is crucial for advancing PAM, ensuring it continues to grow as an effective tool for conservation, biodiversity research, and understanding ocean systems. The global accessibility of standards-driven, metadata-rich and analysis-ready data provides a scalable foundation from which AI/ML models can be trained. It further serves as a model for tackling big data challenges for the broader earth-system science community where AI-driven research is of increasing prevelance^120,121^.

## Data Availability

The data products used in this project are available at the NOAA National Centers for Environmental Information Passive Acoustic Data Archive, https://www.ncei.noaa.gov/products/passive-acoustic-data.

## Acknowledgements

This research was supported by NOAA IOOS, Bureau of Ocean Energy Management, U.S. Navy Living Marine Resources, and Office of Naval Research (award #s: NA21NOS0120100 and N00014-22-1-2601). Brian Stone and Trevor Golden played key roles in developing workflows to access sound levels from three different repositories, integrate acoustic and non-acoustic variables, and the entire *SoundCoop* portal. CP and PyPAM were supported by the Research Foundation Flanders (FWO) as part of the Belgian contribution to the LifeWatch ESFRI, Grant number I002021N. KT and OB were supported by the Helmholtz strategic infrastructure FRAM under grant number AWI-PS100_0. EM and AM were supported by NOAA IOOS / SECOORA. JM-O and the AEON project were funded by the Office of Naval Research under award number N00014-20-1-2312.

