## Appendices for "Big Data, Sound Science, Lasting Impact: a framework for passive acoustic monitoring"

**Appendix I.**

Open-source and freeware processing tools, namely, Making Ambient Noise Trends Accessible (MANTA) and the Python Passive Acoustic analysis tool for passive acoustic Monitoring (PyPAM) were chosen to produce the HMD spectra soundscape metric.

MANTA

MANTA calculates the sound pressure spectral density (PSD) levels in units of 1 $\mu Pa^2$ $Hz^{-1}$ using Welch's Method in Matlab. The Discrete Fourier Transform length is equal to the sample rate, a Hann window of equal length is applied to the data and 50% overlap is used. This results in PSD estimates of mean-square pressure amplitude ($\mu Pa^2$) with a frequency resolution of 1 Hz and temporal resolution of 1 s. The 120 PSD estimates from each 1-minute segment are averaged, and the average spectrum for each minute is further processed to the HMD spectrum as dB re 1 $\mu Pa^2$ $Hz^{-1}$, as defined in Martin et al. (2021a). The MANTA outputs for each day are: (1) CSV of the one-minute HMD results; (2) image of the daily long-term spectral average based on the one-minute HMD results, (3) image of the daily spectral probability density with percentiles, and (4) netCDF containing products 2 and 3, deployment-level metadata entered by the user before processing begins, and the calibration coefficients used to compute the calibrated spectrum levels across each processed frequency band. MANTA is available as a 'desktop' application that requires a Matlab license and specific toolboxes to run, and as a standalone executable that does not have a dependency on Matlab making this toolkit available as freeware.

*Calibration*

Sensor sensitivity is entered using a 'Metadata app' within the MANTA program, another graphic user interface, where a set number of manufacturer-derived calibration curves (e.g., HTI-96min) can be called or custom calibration curves can be built. The latter allows the user to ensure the unique sensor, recorder, and pre-amplifier calibration are incorporated for custom systems or commercial systems not yet available in the manufacturer pull-down options. Both flat and frequency-dependent calibration inputs are available. A graphic of the system calibration that MANTA will apply based on the entered information allows the user to verify the calibration content.

*Processing*

Since MANTA was deemed 'community-ready', data were processed from 8 different recording systems in 15 different recording locations. These datasets provided a diverse and rigorous test for MANTA's ability to intuitively apply calibration across a wide range of scenarios and recorder-specific nuances. Outcomes of the testing included: 1) improved handling of time within and between minutes, particularly when minutes are split across files, and when less than 10 seconds within a minute are dropped; 2) each 60-second (or less) interval constrained to a single minute, ensuring that seconds within each bin do not cross two different minutes; 3) the additional output of how many seconds were processed in each one-minute bin; and 4) the need for a structured netCDF that contains the spectra output alongside deployment-specific metadata to facilitate data sharing. New MANTA versions were produced to address needs 1 to 3 above. The final standalone executable version of MANTA that was tested and used in the *SoundCoop* was v9.6.15 pre-release F though some datasets were processed using versions down to v9.6.11.

PyPAM

PyPAM is a Python package to analyze underwater sound[1]. Python is natively open source and operates from a command line interface. The repository was demonstrated in previous publications but did not initially have a function to produce HMD[2,3]. That function was created following methodology defined in Martin et al.[4,5], and implemented in MANTA, and validated against independently computed HMD spectra produced by the original MATLAB code published in Martin et al.[4].

*Calibration*

PyPAM relies on the use of the pyhydrophone package[6]. Both flat and frequency-dependent calibrations can be entered by the user in different formats. Furthermore, this package provides certain functionalities to automatically extract calibration information. For example, for SoundTrap hydrophones the user-entered hydrophone serial number can be used to extract the manufacturer calibration information. For other instruments calibration data can be read from the audio files header[6].

*PyPAM-based Processing (PBP)*

PyPAM-based Processing (PBP) comprises a suite of Python code that extends PyPAM capabilities to enable scalable production of daily HMD data products in local and cloud computing environments. PBP modules include (1) temporal metadata generation and review, (2) daily HMD processing output directly to netCDF files, and (3) summary plot generation. Temporal metadata are extracted from audio files, or from XML metadata files in the case of SoundTrap hydrophones, and stored in daily json files. Each daily json file contains essential metadata for every audio file that contains data within that day: the Uniform Resource Identifier (URI) to locate the file in local or cloud storage, and file start and end times to map input audio data to output one-minute HMD temporal bins. Global and variable attributes, which are needed to produce complete self-describing daily netCDF output, are stored in text files (YAML format), and some output metadata such as software version are automatically populated in the output netCDF.

*Processing*

A one-year continuous dataset from the MBARI-MARS cabled observatory, sampled at 256 kHz, was processed into HMD using PBP. PBP was extended to support more PAM instruments, including the icListen hydrophone used at the MARS site. The output files consist of daily netCDFs and summary plots of the HMD results. The final version of PyPAM and PBP tested and used within the *SoundCoop* was 0.3.0 and 0.2.0, respectively.

### Standard Outputs

A standards-driven netCDF format was created for the daily HMD files. NetCDF is a self-describing, machine-readable data format that supports access and sharing of array-oriented scientific data[7]. Metadata as well as the HMD spectra are stored in a single file, which facilitates data sharing and development of scalable tools for visualization and analysis within and outside of the original team. Metadata about the recording site and monitoring effort including frequency- and time-dependent data quality are captured in JSON files using either NCEI's PassivePacker, a data packaging tool, or PBP's pre-processing workflow.

The netCDF template was designed following standards outlined by the International Organization for Standardization (IHO), netCDF conventions for earth science data (COARDS), climate and forecast metadata conventions for netCDFs (CF conventions), and Earth Science Information Partners' attribute convention for data discovery (ACDD).

Using the data quality content from PassivePacker's metadata JSON, a data quality matrix the same dimensions as the HMD matrix was created for every daily HMD file. The data quality matrix provides a machine-readable way to mask frequency bins and/or time bins when HMD results are present but not calibrated or otherwise of degraded quality. The data quality matrix followed the IOOS Quality

Assurance / Quality Control of Real Time Oceanographic Data (QARTOD) quality tag numbering where 1 = Good; 2 = Not evaluated/Unknown; 3 = Compromised/Questionable, and 4 = Bad/Unusable (U.S. Integrated Ocean Observing System, 2017). Only data with a quality tag of 1 (Good) were used in this analysis.

MANTA output files were processed into single daily netCDFs according to this standard. PyPAM/PBP natively outputs daily HMD results in this netCDF format. By adhering to the same structured file format, netCDFs of data processed with either software can seamlessly integrate into visualization and analytical tools.

**Appendix II**. The metadata content assigned in the netCDF *global attribute* fields and parameters assigned in the *variable* and *coordinate* fields.

| Name | Type | Definition | ACDD Recommendation Level | Example |
|---|---|---|---|---|
| acknowledgement | Global Attribute | Statement on the project and its funders | Recommended | These products support the Passive Acoustic Monitoring National Cyberinfrastructure (SoundCoop) project funded by the NOAA Integrated Ocean Observatory System, Bureau of Ocean Energy Management, U.S. Navy Living Marine Resources, and Office of Naval Research. |
| comment | Global Attribute | Text version of the deployment's data quality | Recommended | Data quality: Good 2021-01-26T20:00:00 to 2021-03-21T18:20:00 from 20Hz to 24000Hz; Data quality: Compromised 2021-01-26T20:00:00 to 2021-03-21T18:20:00 from 0Hz to 20Hz. SoundTrap 300 has a high pass filter at 20 Hz, rendering sound pressure levels below 20 Hz inaccurate. |
| conventions | Global Attribute | Identification of the convention and, if applicable, version used | Highly Recommended | COARDS, CF-1.6, ACDD-1.3 |
| creator_name; creator_role | Global Attribute | Name of person who created the HMD | Recommended | Timothy Rowell, Sofie Van Parijs, Leila Hatch<br><br>Principal Investigator |
| date_created | Global Attribute | Date the netCDF file was created following ISO 8601 format | Recommended | 2023-11-29 |
| geospatial_bounds | Global Attribute | For stationary data, these are listed as points in decimal degrees. | Recommended | POINT (42.43832 -070.54551) |

| | | | | |
|---|---|---|---|---|
| history | Global Attribute | This statement identifies the software and version used to create the HMD and the version of code used to produce the netCDFs | Recommended | Original hybrid millidecade spectra were produced by Timothy Rowell, Sofie Van Parijs, Leila Hatch. NCEI created a single ISO-compliant netCDF_converters from the four MANTA outputs plus additional metadata from the deployment and overall project. Conversion was done using v.1.1.0 of the NCEI MANTA netCDF converter. |
| id | Global Attribute | * Digital object identifier (DOI) for the project | Recommended | https://doi.org/10.25921/ymf8-5k59 |
| infoURL | Global Attribute | Link to organization website | | https://ncei.noaa.gov |
| institution | Global Attribute | Name of institution | Recommended | NOAA National Centers for Environmental Information |
| instrument | Global Attribute | Recording instrument | Suggested | SoundTrap 300 |
| keywords; keywords_vocabulary | Global Attribute | Keywords for the dataset and associated library. For example, those pulled from the Global Change Master Directory (GCMD) | Highly Recommended | GCMD:oceans, GCMD:ocean acoustics, GCMD:ambient noise, intensity, GCMD:marine environment monitoring, marine habitat, sound intensity level in water, soundscapes; GCMD: GCMD Keywords |
| license | Global Attribute | Creative license | Recommended | CC 4.0 |
| naming_authority | Global Attribute | The organization that provides the id for the dataset | Recommended | NOAA National Centers for Environmental Information |
| product_version | Global Attribute | Version of the file | Suggested | v2 |

| | | | | |
|---|---|---|---|---|
| project | Global Attribute | Project name | Recommended | SanctSound, SoundCoop |
| publisher_email; publisher_name; publisher_type; publisher_url | Global Attribute | Information relevant to the publisher of the data | Recommended |; NOAA National Centers for Environmental Information; institution; https://www.ncei.noaa.gov/products/passive-acoustic-data |
| reference | Global Attribute | References related to the processing standards used | Suggested | Original audio recordings are available open-access: https://www.ncei.noaa.gov/maps/passive-acoustic-data/. Computation of single-sided mean-square sound pressure spectral density with 1 Hz resolution followed ISO 18405 3.1.3.13 (International Standard ISO 18405:2017(E), Underwater Acoustics – Terminology. Geneva: ISO). Hybrid millidecade band processing followed Martin et al. (2021; https://doi.org/10.1121/10.0003324) |
| source | Global Attribute | Information on the processing software | Recommended | Data analysis was performed using the Making Ambient Noise Trends Accessible (MANTA, https://bitbucket.org/CLO-BRP/manta-wiki/wiki/Home, see Miksis-Olds et al., 2021; Martin et al., 2021a,b) standalone software (v9.6.13) to produce hybrid millidecade spectra of sound levels from ocean audio recordings. To efficiently tackle large datasets, MANTA is designed around a parallel-processing Matlab package, Raven-X (Dugan et. al., 2014, 2016, and 2018) that uses ordinary multi-core computers to accelerate processing speeds. MANTA calculates the sound pressure spectral density (PSD) levels in units of 1 µPa^2/Hz using Welch's Method in Matlab. The Discrete Fourier Transform length is equal to the sample rate, a Hann window of equal length is applied to the data and 50% overlap is used. This results in PSD estimates of mean-square pressure |

| | | | | |
|---|---|---|---|---|
| | | | | amplitude (µPa^2) with a frequency resolution of 1 Hz and temporal resolution of 1 second. The 120 PSD estimates from each 1-minute segment were averaged, and the average spectrum for each minute was further processed to a hybrid millidecade (HMD) spectrum as dB re 1 µPa^2/Hz, as defined in Martin et al. (2021b). Hybrid millidecades are an efficient means of storing PSD spectra from high sample rate audio files using 1-Hz values up to 435 Hz, then millidecade wide PSD values up to one half of the sampling rate (Martin et al., 2021b). The MANTA outputs for each day are: (1) CSV of the 1 minute HMD results; (2) image of the daily long-term spectral average based on the 1 minute HMD results, (3) image of the daily spectral probability density with percentiles, and (4) NetCDF containing products 2 and 3 in addition to a deployment-level MANTA Metadata output file containing the associated frequency-dependent calibration data used to compute the calibrated spectrum levels. |
| standard_name_vocabulary | Global Attribute | Vocabulary convention used. For example, CF Standard Name Table v80 | Recommended | CF Standard Name Table v80 |
| summary | Global Attribute | A short statement on the purpose of the processed and methods used | Highly Recommended | To understand natural and anthropogenic sound in the ocean, and to compare underwater soundscapes globally, standard methods of analysis must be applied to passive acoustic monitoring (PAM) data. Methods that balance constrained volume and adequate resolution of acoustic spectra have recently been published (Martin et al., 2021a,b). A community effort |

| | | | | |
|---|---|---|---|---|
| | | | | supported by NOAA, BOEM, U.S. Navy, and ONR was initiated to apply these methods to PAM datasets from around the world. This record represents the hybrid millidecade (HMD) spectra of sound levels derived from calibrated passive acoustic data. Daily HMD at 1 minute resolution were created using standalone MANTA software (v9.6.13). These data were recorded at SB01 between January 26, 2021 and March 21, 2021. |
| time_coverage_duration; time_coverage_resolution | Global Attribute | Format for duration and resolution of the processing output. The example shows a one day file duration of 60 second resolution | Recommended | P1D; P60S |
| time_offset | Global Attribute | Hours from UTC | | 0 hours from UTC |
| title | Global Attribute | Name of the dataset | Highly Recommended | Hybrid Millidecade Band Sound Pressure Levels Computed at 1 Minute Resolution from Oceanic Passive Acoustic Monitoring Recordings at SB01 |
| SamplingRate | Global Attribute | Deployment sample rate in Hz | | 48000 |
| PreampFixedGain_dB; CalibrationFrequency_Hz; CalibrationSensitivity_dB_re_1VperRefPress; CalibrationDate | Global Attribute | Text based details about the hydrophone/recording sensitivity and calibration, if provided | | |
| comment | Variable Attribute | Description of the variable calculation | | Computation of single-sided mean-square sound pressure spectral density followed ISO 18405 3.1.3.13. |

| | | | | |
|---|---|---|---|---|
| coverage_content_type | Variable Attribute | ISO19115-1 code that indicates the source of the data | Highly Recommended | physicalMeasurement |
| long_name | Variable Attribute | Long descriptive name for the variable | Highly Recommended | Single-sided mean-square sound pressure spectral density re 1 micropascal^2/Hz |
| standard_name | Variable Attribute | Long descriptive name taken from controlled vocabulary of variable names | Highly Recommended | sound_intensity_in_water |
| units | Variable & Coordinate Attributes | Units of the variable’s data values | Highly Recommended | dB |

**Appendix III.** Datasets used to demonstrate value from comparative analyses. Details include project name, region where data were recorded, recording site name, latitude (Lat), and longitude (Lon), depth of the instrument (Instr.) and seafloor, start and end of the analysis period, effective frequency bandwidth of the calibrated audio data, how long sound was recorded (duty cycle), type of recording instrument, software used to process the audio data into HMD spectra, and the repository (repo) from where the HMD spectra were accessed. The asterisks by instrument and bottom depth indicate values may vary slightly due to multiple deployments represented within the analysis period. When sites are not recording continuously (Cont.), the recording duration is listed first, followed by the interval between the start of two consecutive recordings. For example, the DSG recorder used in the ESONS project recorded for 2 minutes every hour. MANTA refers to the Making Ambient Noise Trends Accessible freeware processing software; PyPAM refers to the Python passive acoustic analysis tool for Passive Acoustic Monitoring open source processing software; NMS is National Marine Sanctuary; GCP refers to the NCEI Passive Acoustic Data Archive GCP bucket; AWS refers to the MBARI MARS AWS bucket; and Axiom is Axiom Data Science's Research Workspace.

| **Case Study** | Project Name | Region | Site Name | Lat | Lon | Instr. Depth (m)* | Bottom Depth (m)* | Analysis Start | Analysis End | Effective Bandwidth (Hz) | Duty Cycle | Instr. Type | Software | Repo |
|---|---|---|---|---|---|---|---|---|---|---|---|---|---|---|
| **1** | AFSC-ALTIMA | Arctic Ocean<br>Chukchi Plateau | AU_CH01 | 75.10 | -168.00 | 118.6 | 166 | 2008-10-13 | 2009-10-05 | 20-4,096 /<br>20-8,192 | 8 min/20 min | AURAL | MANTA | GCP |
| **2 & 3** | AEON | Gulf of Maine<br>Wilkinson Basin | AEON5 | 42.87 | -70.02 | 133.5 | 135 | 2021-02-14 | 2021-12-31 | 5-7,072 | 45 min/1 hr | AMAR | MANTA | GCP |
| **2** | ESONS | Southeast U.S.<br>May River | 37M | 32.20 | -80.79 | 5.1 | 5.3 | 2021-01-01 | 2021-12-31 | 50-40,000 | 2 min/1 hr | DSG | MANTA | Axiom |
| **2 & 3** | MBARI-MARS | Central California Coast | MARS | 36.71 | -122.19 | 890 | 891 | 2021-01-01 | 2021-12-31 | 10-200,000 | Cont. | icListen | PyPAM | AWS |

| | | | | | | | | | | | | | |
|---|---|---|---|---|---|---|---|---|---|---|---|---|---|
| | | Monterey Bay | | | | | | | | | | | |
| **2** | NEFSC | Gulf of Maine Coastal | Monhegan Island | 43.72 | -69.30 | 58 | 61 | 2021-02-05 | 2021-12-18 | 20-24,000 | Cont. | Sound Trap | MANTA | GCP |
| | | | Petit Manan | 44.32 | -67.86 | 58 | 61 | 2021-01-01 | 2021-12-10 | | Cont. | Sound Trap | | |
| **2 & 3** | NRS | Central California Coast Cordell Bank NMS | NRS11 | 37.88 | -123.44 | 500 | 550 | 2021-01-01 | 2021-10-04 | 10-2000 | Cont. | AUH | MANTA | GCP |
| **2** | SanctSound | Gulf of Maine Stellwagen Bank NMS | SB01 | 42.44 | -70.55 | 47 | 50 | 2021-01-26 | 2021-12-31 | 20-24,000 | Cont. | Sound Trap | MANTA | GCP |
| | | | SB03 | 42.26 | -70.18 | 42 | 45 | 2021-01-26 | 2021-12-31 | | Cont. | Sound Trap | | GCP |
| **4** | BOEM-Cornell | Coastal Mid Atlantic U.S. | BOEM-VA | 36.90 | -75.69 | 20 | 22 | 2016-08-28 | 2016-09-10 | 10-4,000 | Cont. | MARU | MANTA | GCP |
